# Accurate Identification of Motoneuron Discharges from Ultrasound Images Across the Full Muscle Cross-Section

**DOI:** 10.1101/2023.09.29.560220

**Authors:** Emma Lubel, Robin Rohlén, Bruno Grandi Sgambato, Deren Y Barsakcioglu, Jaime Ibáñez, Meng-Xing Tang, Dario Farina

## Abstract

**Objective:** Non-invasive identification of motoneuron (MN) activity is commonly done using (EMG). However, surface EMG (sEMG) signals detect only superficial sources, at less than approximately 10-mm depth. Intramuscular EMG can detect deep sources, but it is limited to sources within a few mm of the detection site. Conversely, ultrasound (US) images have high spatial resolution across the whole muscle cross-section. The activity of MNs can be extracted from US images due to the movements that MN activation generates in the innervated muscle fibers. Current US-based decomposition methods can accurately identify the location and average twitch induced by MN activity. However, they cannot accurately detect MN discharge times.

**Methods:** Here, we present a method based on the convolutive blind source separation of US images to estimate MN discharge times with high accuracy. The method was validated across 10 participants using concomitant sEMG decomposition as the ground truth.

**Results:** 140 unique MN spike trains were identified from US images, with a rate of agreement (RoA) with sEMG decomposition of 87.4 ± 10.3 %. Over 50% of these MN spike trains had a RoA greater than 90%. Furthermore, with US, we identified additional MUs well beyond the sEMG detection volume, at up to >30 mm below the skin.

**Conclusion:** The proposed method can identify discharges of MNs innervating muscle fibers in a large range of depths within the muscle from US images.

**Significance:** The proposed methodology can non-invasively interface with the outer layers of the central nervous system innervating muscles across the full cross-section.

## I. Introduction

Amotor unit (MU) comprises a motoneuron (MN) and the muscle fibers it innervates (the latter referred to as the muscle unit). The MU is the smallest functional element of the muscular system. MUs have been studied within the fields of human neuromechanics and clinical diagnosis, classically relying on electrical recordings of the action potentials (APs) of the muscle unit. Because the neuromuscular junction (NMJ) is a highly reliable synaptic connection, the correspondence between an AP discharged by the innervating MN and those discharged by the innervated fibers is one-to-one [1]. The sum of the electrical activity of MU APs is the electromyogram (EMG), which can be detected either at the surface of the skin (sEMG) or intramuscularly (iEMG). sEMG usually presents a low spatial resolution [2]–[4] and small surface detection volume [4], [5], while iEMG is very selective around the electrode site [6]. In each case, a large proportion of the Mus in the muscle are not accessible.

When the muscle fibers are activated, their membranes are electrically depolarized and shorten because of the formation of actin-myosin cross bridges. The contraction event of the fibers of a MU is called a twitch. Much like the transfer of the electrical signal at the NMJ, this electromechanical coupling is one-to-one, meaning that each mechanical twitch is precisely related to an individual MN discharge instance [7]. Thus, the activity is translated from the electrical domain to the mechanical domain. In theory, a recording of the deformation within the muscle could be processed to identify the precise time of MU twitches and, thus, to detect the neural input received by the muscle from the spinal cord. In contrast to the resolution and penetration issues of EMG, ultrafast ultrasound (US) could, in principle, detect the small movements of the muscle units in any location in the muscle with high temporal resolution.

However, the individual MU velocity twitches are very small (approximately 3 mm/s [7]), and the territories of different MUs overlap in space [8], thus a given muscle region will have a complex deformation field [9]. When the fibers contract, they push and pull on each other and nearby tissues, setting up propagating waves within the region [10], [11]. Furthermore, deformation due to connective tissue, bones, and blood vessels is larger than the MU twitches, and structures mechanically couple, which generate mechanical noise. Separating individual MU activity from these noise sources and from the activity of other MUs in US images is challenging.

Despite the complexity of the problem, methods have been proposed to extract neural information from recordings using ultrafast US imaging to detect muscle deformation [12]–[14]. Current methods are based on blind source separation (BSS) of linear instantaneous mixtures, which optimize sparse spatial filters used as weights for the linear combination of pixels to extract temporal information. Using these approaches, estimates of MU locations have a high repeatability. However, the discharge times are estimated with relatively low accuracy (with an agreement between US and the gold standard EMG of only ∼30% of the discharges) [15]. Spatial linear combinations of pixels (which is equivalent to spatial filtering of the velocity maps) cannot compensate for the long duration of the MU velocity twitch profiles. Instead, an inverse transformation which also contains the time variable is required. Hence, an anti-transformation in space and time may be more appropriate for separating the individual MU activity from that of other MUs and noise.

Because of the electromechanical coupling, the generation of a twitch due to a neural discharge can be modelled as a convolution of a delta function centered at the discharge instant with the twitch response. Therefore, we propose that, compared to an instantaneous model, a convolutive generative model will more effectively account for the specific temporal dynamics of MU activity, allowing for better identification of the times of occurrence of MU discharges and better separation from noise. We hypothesized that solving the unmixing problem under the assumption of a convolutive mixture will solve the decomposition challenge of US deformation fields into individual MU contributions with better accuracy than with instantaneous models.

Here, we propose a method for decomposing US image series into MN activity using a convolutive BSS method. Unlike previous methods for US analysis [12]-[15], our method is completely stand-alone and requires no concurrent EMG recording. We validated the method using simultaneously recorded US and high-density sEMG (HDsEMG) signals at low force levels from 10 participants. The HDsEMG was decoded using a previously validated EMG decomposition algorithm. We used our proposed method to decode the deformation fields calculated from US and quantified its performance using the rate of agreement (RoA) with the HDsEMG decomposition output. This comparison was performed only for relatively superficial MUs due to the limited detection volume of the sEMG. Further, we quantified the time difference between HDsEMG-estimated discharge times and the US-estimated discharge times, as well as the range of this difference across the 30-s recording intervals. For all US-identified sources, we further performed a spike-triggered average (STA) of the sEMG using the estimated US discharge times as triggers. This provided estimated APs for each identified source (not only for the sources commonly detected between US and sEMG). When the STA was significantly above the noise level, the corresponding source was interpreted as the series of discharges of a MU. We show that our proposed method can extract highly accurate neural activity across the full muscle cross-section, with many units detected deeper than the penetration depth of electrical signals. The method could, therefore, pave the way for an alternative interfacing pathway with the output layers of the spinal cord circuitry.

## II. Methods

In this section, we describe the convolutive model and implementation used for decoding the velocity images from US into neural spike trains. We then describe the validation approach.

### A. Model

When MUs are activated, a series of APs travel through the muscle fibers causing a series of twitch responses of the fibers. The AP discharge times can be modelled as a series of delta functions and the twitches as impulse responses. The resulting US images are then the sum of the convolutions of the delta functions and their impulse responses. Additional noise, such as that from the US imaging and velocity tracking, also contributes to generating the US image series.

Let *x*_*i*_*(t)* with *i* = 1, 2, …, *m* denote the axial velocity in the *i*^*th*^ pixel of the US images, evolving over time *t* ∈ [0, *T]*. Then, we can represent *x*_*i*_*(t)* as the result of the following convolutional model:

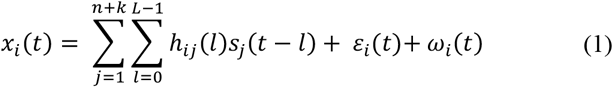

where *n* is the number of MU sources, *k* is the number of non-MU sparse sources (e.g., cyclo-stationary movements of structures within the muscle, such as blood vessel pulsations), *s*_*j*_*(t)* is the *j*^*th*^ source as a function of time *t, h*_*ij*_*(l)* is the impulse response (filter) for the *i*^*th*^ pixel and *j*^*th*^ source, with finite duration *L*, ε_*i*_*(t)* is the additive non-white noise (e.g., bone movement), and ω_*i*_*(t)* is the additive white noise at the *i*^*th*^ pixel. In the isometric contractions considered here, we assumed ε_*i*_*(t)* to be negligible. The *n* sources consist of a series of delta functions, i.e., vectors of ‘0’s and ‘1’s in the time domain. The twitch profile for a specific source, i.e., *h*_*ij*_*(l)*, is assumed to be the same for each discharge.

Eq. (1) has the same form as the generation model of EMG signals [16]. Nonetheless, velocity twitches in US have a much longer time support than APs in EMG, and usually, the velocity twitches are longer than the average duration of the inter-spike interval (ISI) [7], [9]. Moreover, the signal-to-noise ratio (SNR) of single-unit velocity twitches is much lower than that of single-unit APs. The identification of the MU discharges from the US deformation maps can, therefore, be approached partly with methods developed for surface EMG decomposition but with adaptations due to the more challenging problem. In the following, we present a BSS approach for MU discharge times from US images partly derived from previous EMG processing methods [16], [17], but adapted to the US signal properties.

The velocity signal in Eq. (1) corresponds to the *i*^*th*^ pixel of the recording. By extending Eq. (1) to all pixels, we obtain the axial velocities in matrix form ***x****(t)* = *[x*_*1*_*(t), x*_2_*(t)*, …, *x*_*m*_*(t)]*, where *m* is the number of pixels. The *n* + *k* sources can also be represented in matrix form as *s(t)* = *[s*_*1*_*(t), s*_2_*(t)*, …, *s*_*n*+*k*_*(t)]*.

### B. Implementation

To identify the *n* + *k* sparse sources in *s(t)* in Equation (1), the problem is reformulated to a standard linear independent component analysis (ICA) model [18], which is an effective model in identifying sparse components [19]. Given the finite duration of the filters (*L*), the convolutive mixture can be written as a linear instantaneous mixture of a new set of sources, which contains the original sources and their delayed versions. The delay of the sources is used to write convolutions as matrix multiplications so that each source is delayed *L* times, from *1* to *L* samples (i.e., the duration of the impulse response). We can further delay the velocity signals to maintain a large proportion of observations with respect to sources. Assuming the sources are delayed by up to *L* samples and the observations by up to *R* samples, we obtain the following new vectors:

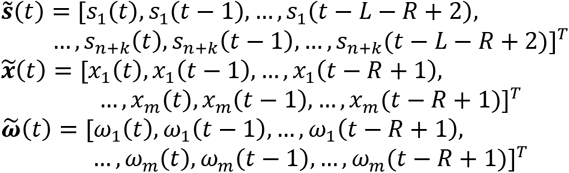

With the above definitions, and setting ε_*i*_*(t)* to zero, we can define the following equivalent linear instantaneous model:

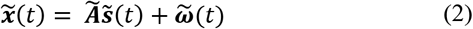

This can now be solved using standard ICA methods and used to estimate *s(t)*.

Given a series of US deformation maps, our first processing step is to divide the data into sub-grids (of size 10 by 10 pixels, corresponding to 3 by 3 mm) such that all subsequent processing is done on a reduced area of the map. The reason for this segmentation is two-fold: firstly, this reduces the number of sources expected in the region and thus simplifies the separation; secondly, given that the next step is to extend the observations by a factor of *R*, we limit the number of observations to reduce the size of the extended matrix and thereby the computational load. Once the processing is complete on one sub-grid of the US image sequence, we slide the grid by 5 pixels (resulting in a partial overlap of 5 pixels between one window of data and the next window) and repeat the processing for the new data. This is equivalent to limiting the number of observations *m* in Eq. (1) to 100 pixels for each decomposition stage.

Once the data has been windowed, vectorized into 2D and standardized such that each row has mean 0 and standard deviation 1, we extend the data by introducing *R* delayed versions of each observation. Following this, the mean is subtracted along each dimension. Next, we whiten the extended observation matrix 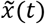, such that they are uncorrelated for time lag zero and have unit variance. To whiten, we use eigenvalue decomposition of the covariance matrix of 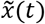, which is a square matrix with size *m×R* by *m×R*. The theoretical number of non-zero eigenvalues is, however, *(n+k)×L<m×R* because the observations are linear combinations of *(n+k)×L* sources. Therefore, the covariance matrix of 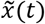 does not have full rank and has *(m×R)((n+k)×L)* non-zero eigenvalues. Because *n+k* is not known a priori and because the estimated eigenvalues are in practice all different from zero due to noise, the number of eigenvalues retained for source estimation has to be blindly estimated. In practice, we apply a threshold on the eigenvalues, and assign those eigenvalues smaller than the threshold to noise and the ones remaining to the sources [20]. We choose a threshold of 70% empirically, however this could be adapted participant-wise (see Discussion). By employing this thresholding step, we implicitly assume that the additional noise is a stationary, temporally and spatially white, zero-mean random process. This is explained in detail in step 5 of the pseudocode in section II.C and is a crucial step for noise removal.

A key assumption for the whitening is that the autocorrelation matrix for time lag zero of the extended sources is diagonal. This property is not valid for generic delayed sources. In our case, because the sources are series of delta functions, the assumption holds as long as the maximum delay *L+R* is shorter than all the ISIs [16], [17]. While this is the case for the mathematically similar problem of EMG decomposition [2], [16], [17], it is not valid for US, where the velocity twitches have a large time support [7], [9], [21]. Therefore, when the sources are extended, there will be one or more delayed source with at least one discharge in common with the original source and potentially other discharges in common with other delayed versions of the same source. In this situation, the scalar product between extended sources at the time lag zero is not always zero, and the autocovariance matrix of the extended sources is not exactly diagonal. Nonetheless, we notice that the effect of non-zero values outside the diagonal of the covariance matrix of extended sources should be negligible due to the small number of nonzero entries with respect to the total number of off-diagonal elements. This is a direct consequence of the sparsity of the sources. Thus, the whitening is minimally affected by impulse responses longer than the ISI, as we later verified empirically. This practically means that we can deconvolve the sources and the unknown filters in the case of sparse sources even if the filter impulse responses are longer than the average distance between activation instants.

After whitening, we applied a fixed-point algorithm to estimate the separation vector and thus separate the sources, with additional Gram-Schmidt orthogonalization to ensure orthonormality and to increase the number of identified sources [22]. Given the high levels of noise within the data, the filters estimated through the fixed-point iteration are unable to separate the source signals and noise fully. In practice, the estimated sources are not ideal trains of delta functions, and an advanced peak detection approach is needed. Hence, we applied a further blind deconvolution step on each estimated source to identify the MU spike train times. Here, the estimated source was extended and delayed, and a second fixed-point algorithm was used to estimate the deconvolved signal, as suggested in a previous work that used linear instantaneous mixing models for ultrafast US decomposition [21]. Finally, the peaks of the estimated sources were identified using a peak-finding algorithm based on the height of the peaks and the time between them [21].

The aim of each of the fixed-point algorithms is to maximize the sparsity of the source vector. This is done using a contrast function G(x) in the form of its first derivative g(x) and its second derivative g’(x). For the first fixed point algorithm (pseudocode step 7.2.1), we used *G(x)* = log (cosh(x)) to maximize the kurtosis of the estimated signals. For the blind deconvolution peak finding algorithm, we used 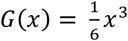, thus maximizing the skewness of the estimated signal.

Following the second fixed-point algorithm, the identified peaks were used to calculate the variability in the predicted discharge times under the assumption that an accurate MU spike train will present some level of regularity in the discharge times. The process was then continued iteratively whilst the variability of discharge times decreased. As such, the iterative process continued to improve the source estimation until it no longer met an improvement threshold. Then, the estimated source was retained if it fulfilled the criteria set for a MU source. The whole process was then repeated for a set *N*_*iter*_ times before the window was moved and the process repeated.

Given the k non-MU sparse sources in the convolution term in Equation (1), some estimated sources reflected non-MU activity, such as blood vessel pulsations. An example of one such output is shown in the Supplementary Material (i). We then selected putative MUs based on the expected features of a sequence of MU discharges – the discharge variability should be low, and the energy of the signal within the expected range for MU discharge (6 - 16 Hz used for low force level contractions) should be high. Thresholds for these cut-offs were chosen empirically: for any given unit if the mean absolute deviation in the discharge time was greater than 25 ms, or if less than 20% of the energy was in the 6 - 14 Hz band (the energy band expected for low contraction level MU activity), the estimated source was disregarded.

Finally, neighboring decomposition windows from the full velocity map often detect MUs in common, given that MU territories are often larger than 3 by 3 mm in the tibialis anterior (TA) [23]. To detect common MUs among decompositions, we calculated the cross-correlation between estimated sources. If the cross correlation was greater than 30%, the sources were considered estimates of the discharge series of the same unit [24]. In this case, the estimated source with the lowest discharge variability was retained, and the others discarded.

Through the above process, we obtain a two-dimensional map of the muscle with localizations of estimated MUs within the muscle cross-section, alongside an estimation of their discharge times.

### C. Algorithm

The process described in section II.A is presented in pseudocode in Table 1. The blind deconvolution peak finding algorithm is presented elsewhere [21].

**TABLE 1.**
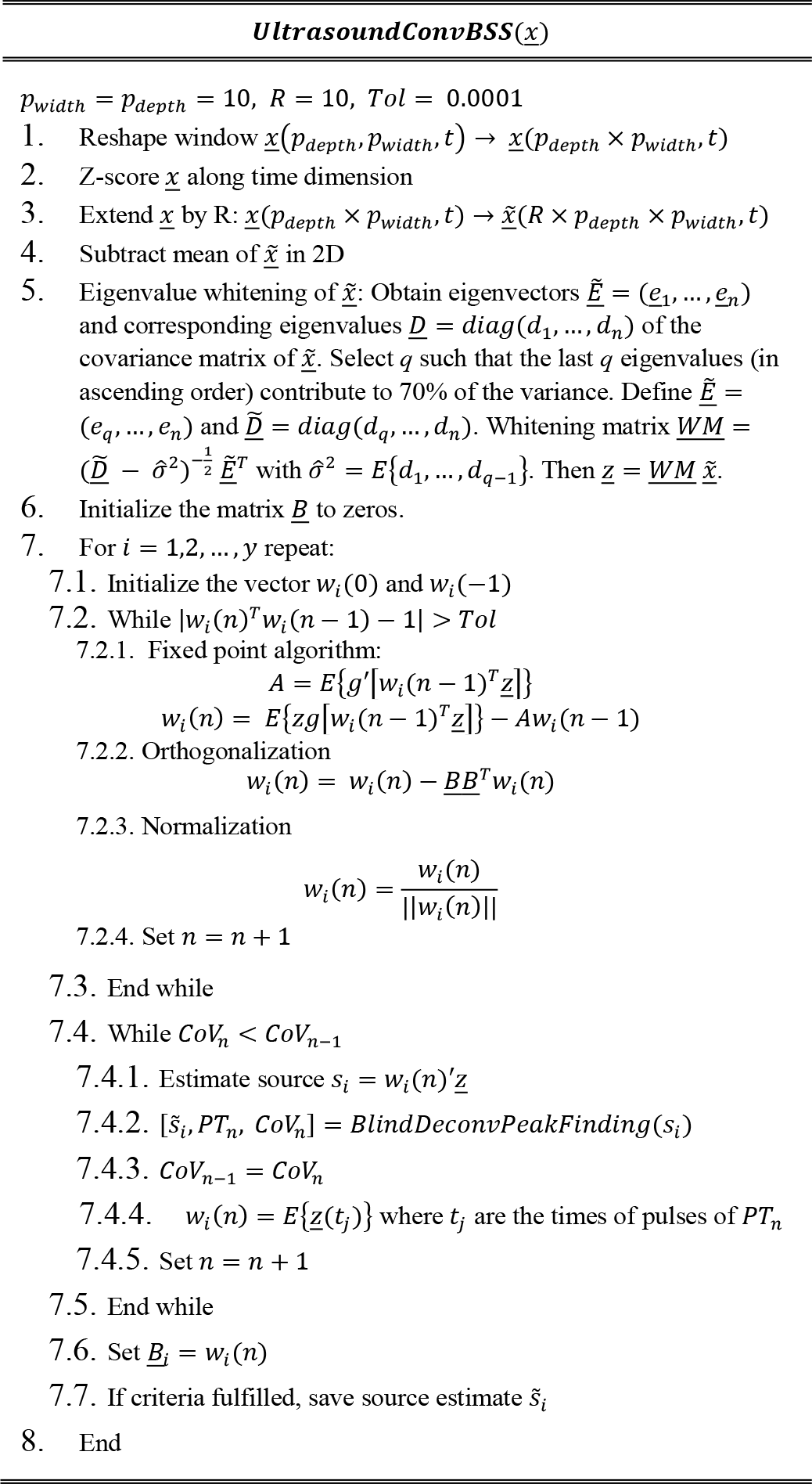
Pseudocode.

### D. Experimental validation

For the validation of the proposed method, we used experimental data previously collected [9] (10 participants, 26.2 ± 2.9 years, 173.2 ± 7.3 cm, 69.1 ± 12.0 kg). In brief, HDsEMG and US data were concurrently recorded during low-force isometric contractions with online feedback on the activity of a small number of MUs online decomposed from surface EMG. This section provides details on the experimental setup, data processing, and data analysis.

### 1) Ethical approval

The experiments were approved by the Imperial College Research Ethics Committee (reference: 20IC6422) in accordance with the Declaration of Helsinki. Before data collection, the volunteers were briefed on the study, presented with a participant information form, allowed to ask any questions, and asked to sign a consent form. As the datasets acquired are large, the data were not registered in a public database but may be made available, as appropriate, upon request.

### 2) Equipment and synchronization

The US data were acquired using the Vantage Research Ultrasound Platform (Verasonics Vantage 256, Kirkland, WA, USA) using an L11-4v transducer with 128 elements and a center frequency of 7.24 MHz. A recording time of 30 s was used, at a frame rate of 1000 Hz due to plane wave imaging with a single angle. Hence, 30,000 frames (357 by 128 pixels) of US data were recorded per trial, followed by delay and sum beamforming. Two HDsEMG grids were used (64 channels; 5 columns and 13 rows; gold coated; 8 mm interelectrode distance; OT Bioelettronica, Torino, Italy). The signals were recorded in monopolar derivation, amplified, sampled at 2048 Hz, A/D converted to 16 bits with gain 150, and digitally bandpass filtered (10–500 Hz), using an EMG pre-amp and a Quattrocento Amplifier (OT Bioelettronica, Torino, Italy). The systems were synchronized using a 1 *μ*s active low output from the Verasonics system, elongated in time using an Arduino UNO and fed into the EMG amplifier.

### 3) Experimental set-up

The TA muscle was used for this experiment (for reasoning, see [9]). The skin over the TA was shaved and cleansed with a chemical abrasive and alcohol. Next, two HDsEMG grids were placed along the length of the muscle fibers, leaving an approximately 1.5 cm gap between them, roughly in the middle of the length of the muscle. The grids were secured with medical tape and Tegaderm Film Dressings. The US probe was placed perpendicular to the length of the muscle in the gap between the EMG grids, secured using a custom-built probe holder and a water-based gel was applied between the probe and the skin. It is widely assumed that due to the isovolumetric nature of muscle fibers [25]–[28], the muscle fiber shortening will result in a cross-sectional increase in area [29], [30], which has been shown experimentally [31], [32]. Thus, we measured local deformations perpendicular to the length of the muscle as a proxy for muscle shortening. We oriented the probe this way to increase the number of Mus crossing the US imaging plane, thus increasing the number of units detectable. The participant was seated comfortably, and their leg was secured in an ankle dynamometer to record force. A screen was placed such that the participant could see it to be used for feedback.

### 4) Experimental protocol

Using the methodology presented in [33], real-time decomposition of the HDsEMG signal was performed, and visual feedback in the form of MU discharges was provided to the participant. Training for the decomposition was done on a signal recorded at constant force corresponding to 10 % of the maximum voluntary contraction force. Following training, the participant recruited an individual MU and held a constant force, at which point the experimenter began a 30-s US recording. After a 60-s rest period, the participant repeated this contraction. This was repeated with an increasing number of concurrently active MUs, up to a total of 6 (or fewer, if fewer were decomposed).

### 5) Processing and decomposition

The HDsEMG data were re-processed using extensively validated offline decomposition methods [16] to ensure high decomposition accuracy. Following this, the estimated MU discharges were manually refined and edited by an expert [34].

A 2D autocorrelation method [35] was used on the reconstructed radio-frequency US data to calculate the tissue velocity, using a sliding window of 2 ms in time and 1 mm in depth. A negative velocity indicated motion away from the probe (and hence the skin). Then, the velocity data were processed, as described in sections II A-C. The outputs of this processing were the estimated MU discharge times.

### 6) Data analysis

For each MU spike train identified using US, the depth of the 3 by 3 mm window in which the unit was identified was recorded. For each MU, the firing rate, coefficient of variation of the ISI (CoV, defined as the standard deviation divided by the mean of the ISI), and the pulse-to-noise ratio (PNR, defined as the average peak height divided by the average baseline noise) was calculated. For the validation, MU spike trains identified from both EMG and US had to be paired. To do so, the number of matched discharges between the two modalities was calculated within a window of 0-5 ms. If more than 30% of all discharges were matched, the units were defined as common units [24]. The RoA was calculated for each EMG-US unit pair as 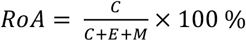, with *C* = number of correct spikes, *E* = number of extra spikes, *M* = number of missed spikes. Furthermore, the average time difference between the EMG and US-identified spikes was calculated, as well as its standard deviation over the 30-s recording.

For each US-identified MU spike train (including the MUs not identified by sEMG decomposition), we spike-triggered averaged the raw monopolar EMG signals using the US-estimated discharges as trigger. This was done to verify that the sources identified from US and not from sEMG were MU activities. The rationale was that if the estimated sources were MU activity, then the STA of the sEMG from these sources should have resulted in waveforms above the noise level. Therefore, the SNR of the triggered sources was calculated as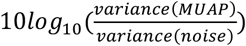.

Descriptive statistics are provided for results, including mean, median, and standard deviation. The characteristics of the population of MUs identified by sEMG and US were compared using a t-tests, given the large sample sizes. The significance level was set as p < 0.05. The p-values are stated on each graph with 3 significant figures, and any p-values lower than 0.001 are labelled as p < 0.001.

## III. Results

Fig. 1 shows an example of the processing steps for an experimental signal, exemplifying the steps of the method, including extending, whitening, and the fixed-point iteration. Across 10 participants, 641 unique MU spike trains were identified using HDsEMG and 409 using US. Of these, 140 were determined to be matched between the two modalities.

**Fig. 1.**
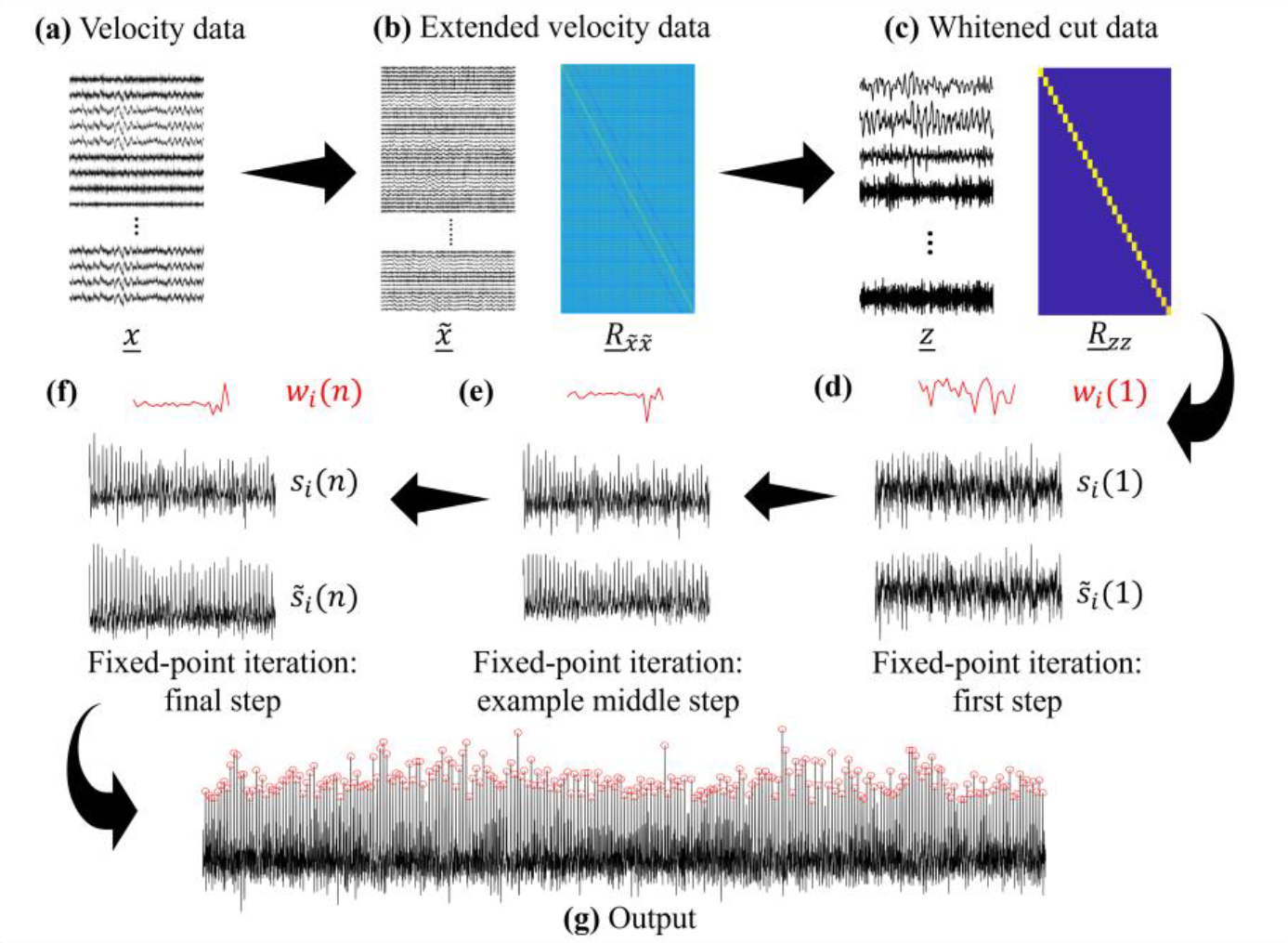
Illustration of the proposed method to detect motor unit (MU) discharge times from ultrasound (US). A 2.5-s interval of data is shown for (a)-(f) for clarity. (g) shows the full output over the 30-s recording duration. (a) Input velocity data with dimensions *pixels × time*. (b) Extended velocity data and autocorrelation matrix of the extended velocity data (*<u>R</u>*_*xx*_). (c) Whitened data after removal of noise components and autocorrelation matrix of whitened data (*<u>R</u>*_*zz*_), showing approximate diagonality. (d) First step of the fixed-point iteration, showing the projection vector *w*_*i*_*(1)* in red (randomly initialized), the estimated source *s*_*i*_(1) and the estimated source after the integrated blind deconvolution 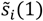. (e) An example of the middle step of the fixed-point iteration. (f) The final step of the fixed-point iteration. The iteration ends here as the coefficient of variation of the estimated discharge times stops decreasing. (g) The full final output of the method shows estimated discharge times as red circles.

Throughout the results, the 140 matched US spike trains will be referred to as the common units. These were used to validate the proposed method by means of comparison with their matched EMG counterparts. In Fig. 2 two 4-s intervals of US source estimation outputs are shown. In the first example (a), no paired EMG unit had been identified. In contrast, the second example (b) is that of a common unit, and the EMG-identified discharge times are shown.

**Fig. 2.**
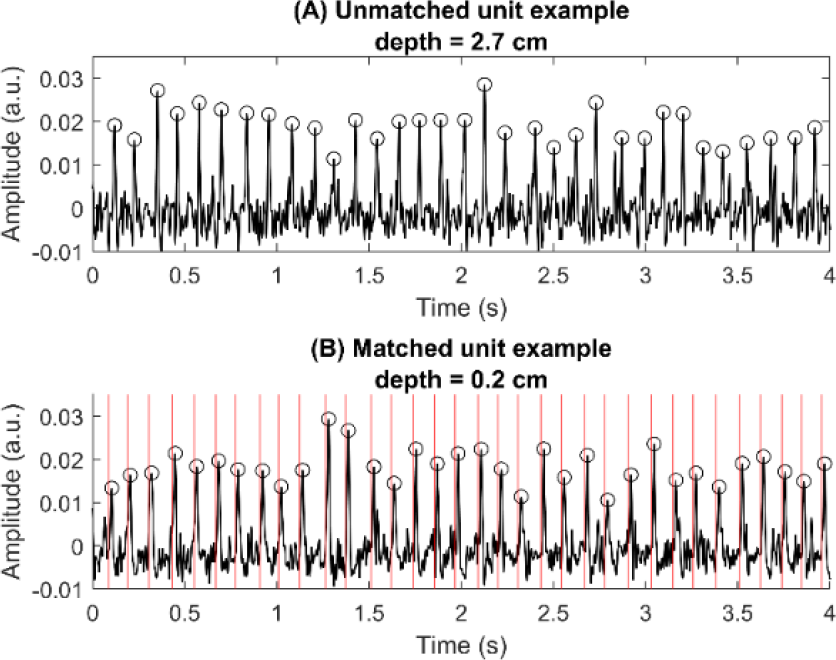
Outputs of the decomposition function applied to the ultrasound (US) image series. In each case, a 4-s interval of the 30-s recording is shown for clarity. Top: The estimated source for a deep motor unit (MU), with identified MU discharge times marked by circles. Bottom: The estimated source for a superficial MU, alongside estimated discharge times for the same MU from the high-density surface electromyography (HDsEMG) decomposition (red lines). Good correspondence is seen between HDsEMG and US-detected discharge times. Furthermore, similar properties are seen for the superficial and deep units.

For the common units, the median RoA (a compound measurement of true positives, false positives, and false negatives) was 90.3 % (mean 87.4 ± 10.3 %), and the median percentage of EMG discharges accurately identified with US (true positives) was 94.9 % (mean 93.1 ± 6.4 %). The distributions of these measures are shown in Fig. 3 (a). Of the 140 units, 51% had RoA > 90%, and 6 units had RoA 100%. The mean firing rate for all common units was 8.5 Hz, therefore on average each MU had 255 firings per recording.

**Fig. 3.**
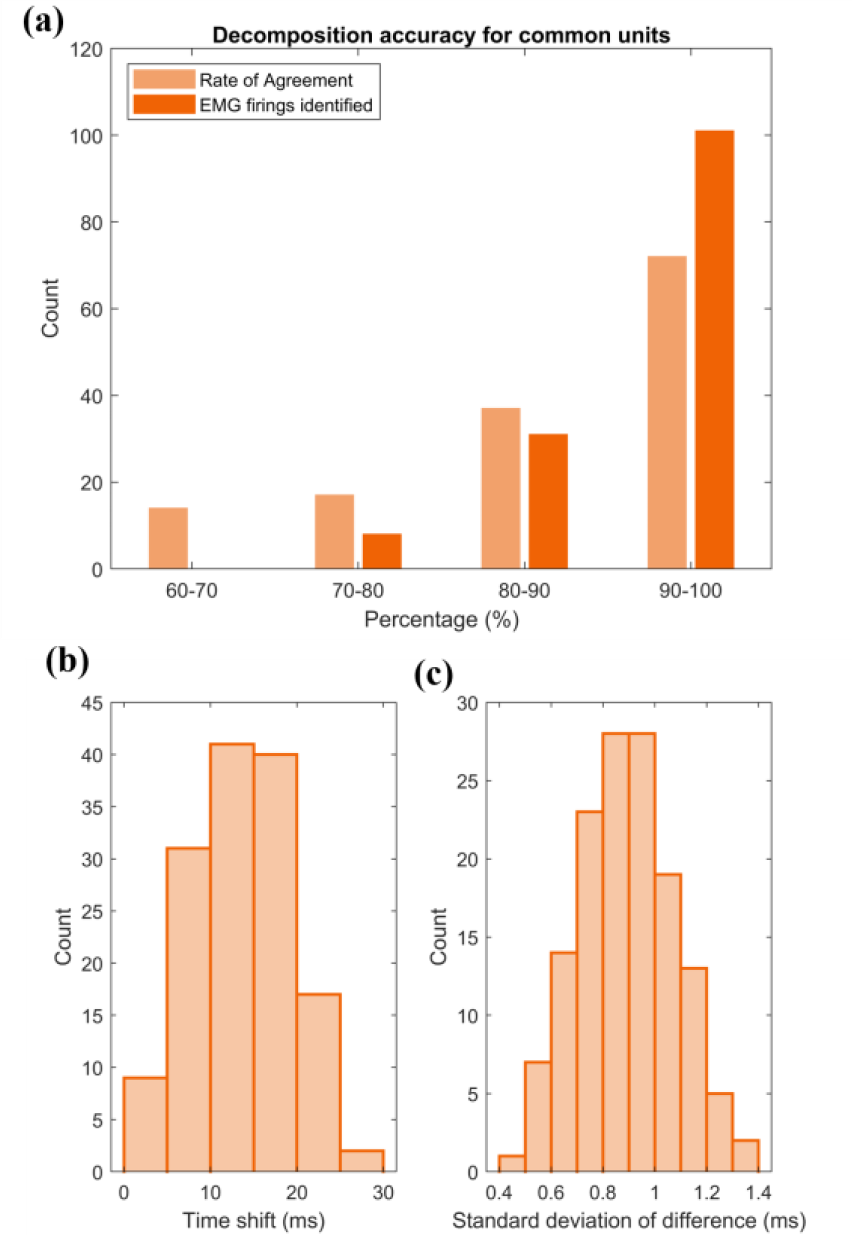
(a) Decomposition accuracy for commonly identified units with respect to the rate of agreement (median 90.3 %) and the percentage of discharges detected using electromyography (EMG), which were also detected using ultrasound (median 94.9 %). (b) The distribution of the time shifts between the discharge times estimated from ultrasound and EMG (mean 13.5 ± 5.4 ms). (c) The distribution of the average variation of this difference (mean 0.89 ± 0.19 ms). It is clear that whilst there is an offset between the identified discharge times from the two modalities, the variation of this offset over the 30-s recording is low, so the offset is relatively constant.

Fig. 3 (b) shows the distributions of the time shifts applied to the US decomposed units to match EMG times, i.e., the delay between the EMG and US-identified discharge times for each unit, and (c) shows the standard deviation of the difference between the discharge times. These plots together show that although, for a given unit, there is a distinct time shift between the EMG and US discharge times (13.5 ± 5.4 ms), the shift is relatively constant across the 30-s period (with a low variability of 0.89 ± 0.19 ms). Thus, the time shift is an expected fixed delay between the electrical and mechanical activity.

Whilst the 140 units identified by both US and EMG have been validated by means of RoA, the remaining 296 MUs identified by US could not be validated with direct comparison with surface EMG (they were outside the detection volume of sEMG). To validate these sources as well, we first analyzed the properties of the 296 remaining units and compared them with those of the common units. In addition, we analyzed the STA of the non-common units, as described later. The distributions of the CoV, discharge rate, and PNR of these two groups are presented Fig. 4. Whilst there was a statistical difference in the CoV (21.6 ± 5.7 and 23.4 ± 5.1 % respectively for common units and other units, p = 0.002) and the PNR (23.0 ± 2.1 and 21.9 ± 1.7 dB respectively for common units and other units, p < 0.001) of the two groups, there was no statistical difference in the discharge rate between the two groups (8.5 ± 0.8 and 8.5 ± 0.9 Hz respectively for common units and other units, p = 0.627). Given that the experimental protocol ensured stable recruitment of EMG-detectable units, it is likely that this slightly reduced the CoV of the discharge times for the EMG-detectable units.

**Fig. 4.**
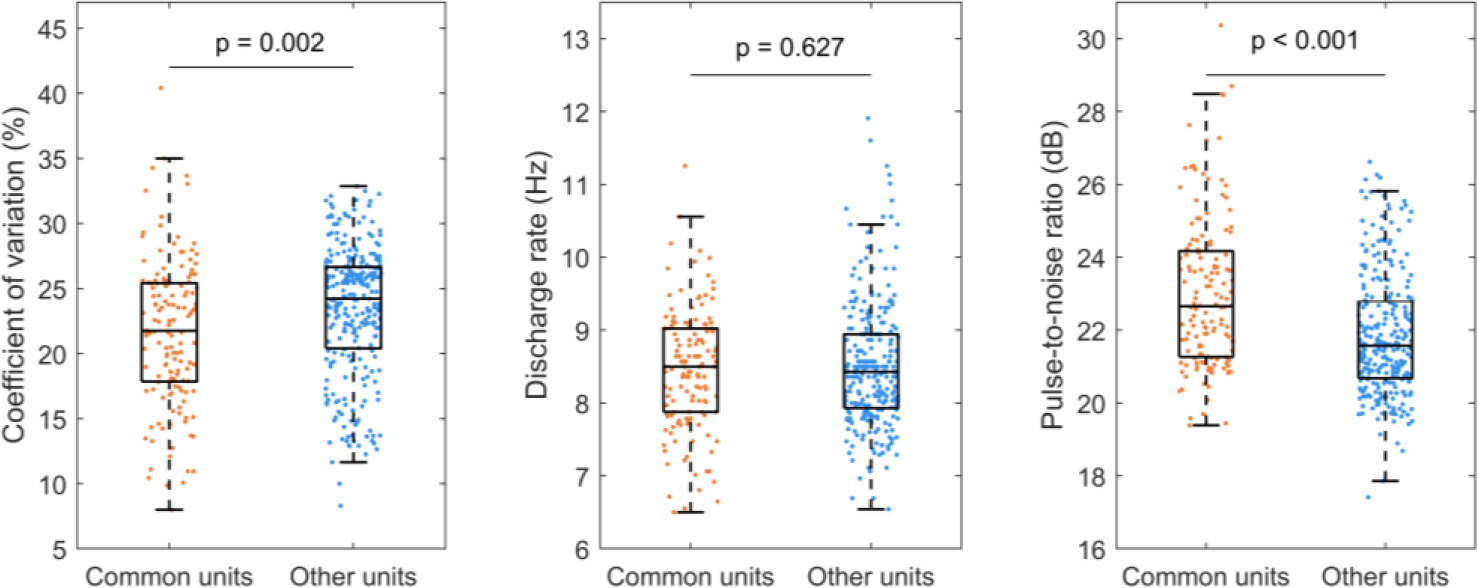
Comparison of common units (those identified by both ultrasound (US) and electromyography (EMG)), n = 140 and other units (those identified only by US, n = 269) based on three metrics: the coefficient of variation, discarge rate, and pulse to noise ratio.

Fig. 5 shows the depth-wise comparison for the two groups of US units (common and non-common with EMG), showing a marked difference in the distribution of the EMG-detectable MUs with respect to the non-EMG-detectable MUs. The great majority of the MUs detected from US but not from EMG were located deeper in the muscle than the common units. The mean depth of the common MUs (3.9 ± 2.7 mm, which corresponds to the detection volume of the EMG) was significantly smaller than the mean depth of the MUs detected only by US (9.8 ± 5.6 mm) (p<0.001). Of the units detected by US alone, 38% were in the most superficial 1 cm of tissue – the expected detection volume of sEMG.

**Fig. 5.**
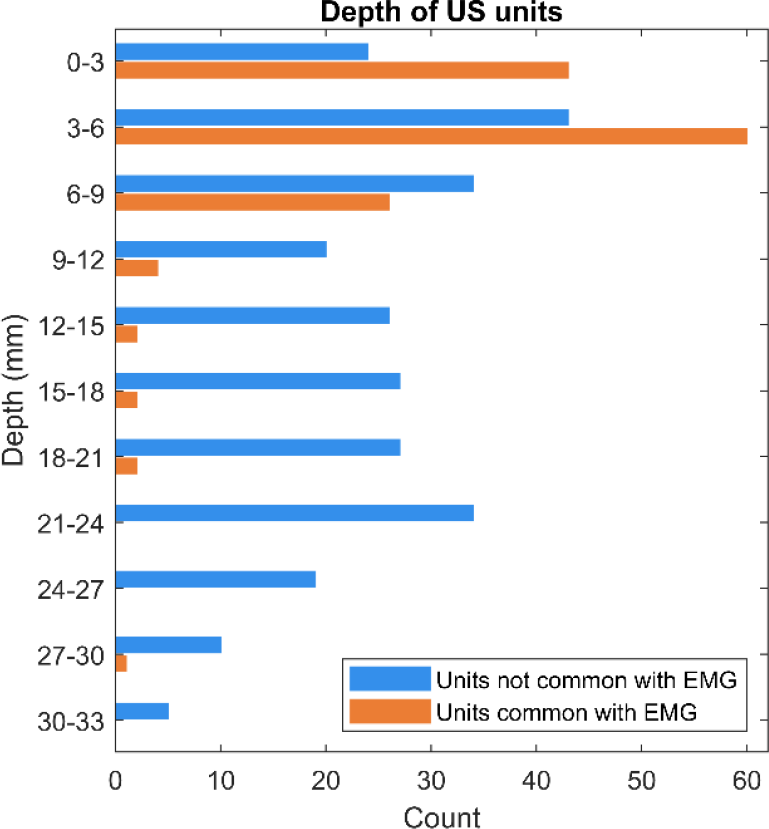
Distribution of motor units (MUs) identified using ultrasound (US) decomposition with respect to depth within the muscle, split between units found in common with surface electromyography (EMG) decomposition (orange, n = 140) and the units that are only identified with the US-based method (blue, n = 269). As expected, the orange distribution peaks within the superficial region of the muscle, as this is where EMG-detectable units lie. Despite the limited detection volume of EMG, peaks are seen in the distribution below 10 mm. These are attributed to split territories and muscle fiber pennation angle. In contrast, the blue distribution spans the full US imaging depth, showing that US can detect deeper MUs than EMG. Further, 38% of MUs detected by only US were in the most superficial 1cm of muscle, therefore US was able to detect units within the EMG detection volume which were not detected using EMG.

Fig. 6 illustrates the estimated waveforms from the STA of sEMG that used US spike trains as triggers, for different depths within the muscle cross-section. For more superficial units (e.g., at a depth of 3 mm), the MUAP peak-to-peak amplitude was large (approx. 0.1 mV) because of the short distance of the fibres from the skin. Conversely, for deep units (e.g., at a depth of 31.5 mm), the MUAP amplitude was small (approx. 0.03 mV) because of the greater distance from the skin surface. However, the classic MUAP waveform shape is evident in all cases and in all cases the peak-to-peak value of this waveform is well above the noise level. Therefore, these sources, while being only detectable from US because too small in the EMG recording, clearly corresponded to MU activity.

**Fig. 6.**
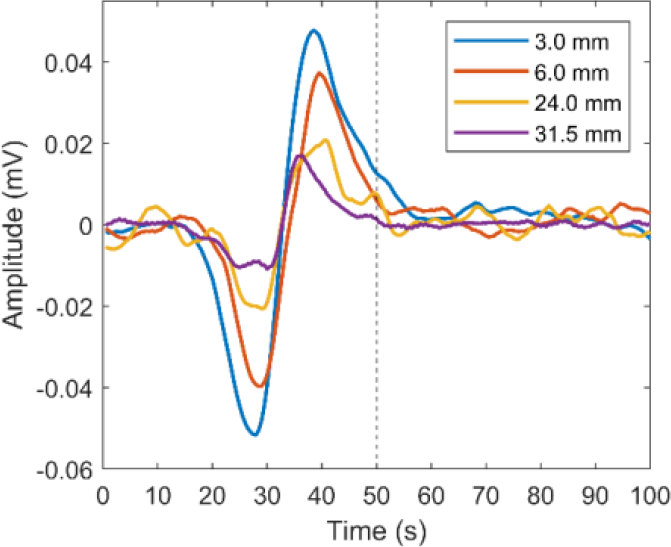
Examples of motor unit action potentials (MUAPs) obtained by performing a spike-triggered average on the electromyography (EMG) signals using discharge times for ultrasound (US) detected MUs. The black dotted line shows the discharge time identified by the US-based method. Superficial MUs show a larger amplitude MUAP with a greater signal-to-noise ratio. However, although the deep units are too distant to be detectable by EMG, the MUAP shape is still visible.

Fig. 7 (b) shows, for all 409 MUs detected from US, the MUAP signal-to-noise ratio as a function of the depth of the unit within the muscle, showing a decrease with depth as the signal gets more attenuated. The average SNR in the most superficial 1 cm of tissue (the approximate detection volume of sEMG) was 29.6 ± 13.3, whereas the average SNR below the most superficial 1 cm of tissue was 15.3 ± 10.8. In contrast, the PNR of the US estimated sources (shown in Fig. 7 (a)) was the same in each region (22.4 ± 1.9 and 22.0 ± 1.9).

**Fig. 7.**
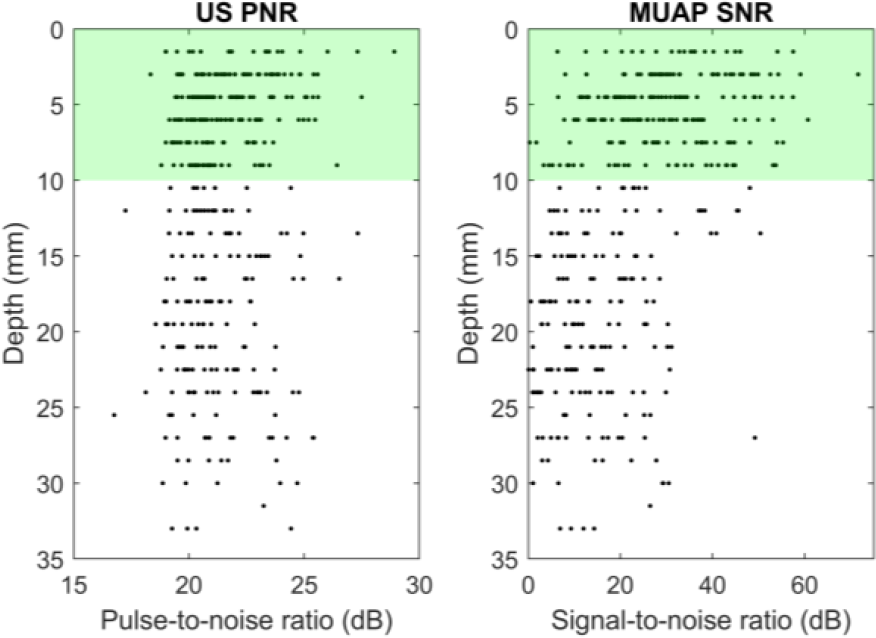
(a) Ultrasound output pulse-to-noise ratio (PNR) as a function of depth. The green region represents the approximate detection volume of EMG. The mean PNR in the green region is 22.4 ± 1.9 (n = 230) and the mean PNR in the white region is 22.0 ± 1.9 (n = 179). Therefore, no degradation of PNR is seen with depth. (b) Distribution of motor unit action potential (MUAP) signal-to-noise ratio (SNR) obtained by spike-triggered averaging of the electromyography (EMG) signals using ultrasound (US) discharge times for all US-detected MUs as a function of depth. In the green region the mean SNR is 29.6 ± 13.3 (n = 230) and in the white region the mean SNR is 15.3 ± 10.8 (n = 179). As the MUs get deeper, the signals get more attenuated, and the signal-to-noise ratio decreases.

## IV. DISCUSSION

We have presented an US decomposition algorithm for estimating MU discharge times in voluntary contractions using convolutive BSS with an integrated blind deconvolution step. We theorized that this technique would separate the MU-related signals from the large noise more successfully than previous methods for US decomposition that rely on linear instantaneous mixture models [12], [15]. Furthermore, by combining traditional convolutional BSS methods with an integrated blind deconvolution, we achieved better signal separation for the noisy input than the convolutive BSS alone. Our method provides a way to non-invasively analyze MN activity for MUs spanning a full muscle cross-section, enabling a more complete way to interface with the central nervous system than has previously been possible.

Across 10 participants, we obtained 409 estimated discharge patterns, 140 of which were matched with corresponding EMG decomposed units. For these 140 units, the median RoA for predicted discharge times was >90 %. Using the RoA as a measure of accuracy provides a conservative result, which accounts for errors in the US decomposition but also in the EMG decomposition and editing. Thus, the accuracy in US decomposition was considerably high.

We validated 140 of the 409 units detected by US using concomitant EMG-based decomposition which has been widely validated in previous works [16], [24], [36]. For the 269 remaining units, we had no ground truth for discharge times since they could not be blindly identified using EMG recordings. However, from the STA of the EMG signals using the US-identified spike trains as triggers, we showed that all 409 estimated sources corresponded to clear MUAP waveforms in the EMG signal, with amplitude above the noise level. Therefore, we argue that all 409 US-identified sources corresponded to MUs. The 269 units identified by US only also showed similar distributions for discharge rates, PNR, and CoV of discharge times to those common with EMG.

In our model, we made some key assumptions of note. One assumption is that the twitch response of a given unit (as recorded by a given pixel) is constant for the duration of the recording. There is evidence to suggest that this may not hold – in force-based experiments, the shapes of MU *force* twitches evolve over time with continuous activation [37], [38]. Although there is a complex relation between the twitch velocity within the muscle volume and the output muscle force due to force transfer throughout the muscle and the tendon, this may indicate that the twitch velocity profile may vary over time, too. Furthermore, previous work has shown that MU twitch profiles sum non-linearly [39], such that the twitch response of a unit depends on the activity of nearby units. This, too, would imply changes of the twitch shape across the recording duration. In summary, changes in twitch shape throughout the recording duration may affect the heights of peaks in the US output. However, at the low contraction levels considered in this study, the influence of this effect was moderate.

With respect to the depth of the units, we conclude that, contrary to EMG, the MU depth is accurately represented in the US decomposition. Of the 140 common units, 92% were located in the most superficial 1 cm of muscle, which is consistent with the low penetration depth of surface EMG. Whilst the further 8% were deeper within the muscle, these units are likely deeper at the location of the US plane than the location where they were detected by the HDsEMG grid due to the pennation angle, or the units have split territories [9], [23], [40]. In the case of split territories, the unit will appear in the decomposition of multiple sub-grids, but only the least noisy output is retained when duplicates are removed. Hence, although a unit appears deep, electrical activity in a more superficial part of the unit’s territory may be detectable using EMG sensors. Example analysis of two such units is shown in the supplementary material (ii).

Furthermore, the US-based method provides insight into units inaccessible using EMG. Fig. 5 shows that MUs are found up to 3 cm into the muscle. The shape of the distribution has some key features, explored in Fig. 8, where the total number of US MUs (409) is considered. From this plot, we observe that most units (56 %) were located in the most superficial 1 cm of the muscle. A potential reason for this observation is that the experiment used online EMG decomposition feedback, thus ensuring that EMG-accessible (hence superficial) units were recruited stably across the 30 seconds. After the most superficial 1 cm, we see a dip in the number of identified MUs, which coincides approximately with the central aponeurosis of the TA muscle. Therefore, the connective tissue will likely limit the motion here, reducing the space for MU identification. At the deepest points, the distribution trails off as participants with smaller muscles do not have muscle below the most superficial 2.5 cm. In summary, although it was possible to identify more MUs from EMG than from US (641 vs 409), likely due to the large muscle area covered by HDsEMG sensors and the bias in the experimental setup due to the EMG online feedback, those MUs identified from US had muscle units distributed across the entire muscle cross-section, contrary to EMG.

**Fig. 8.**
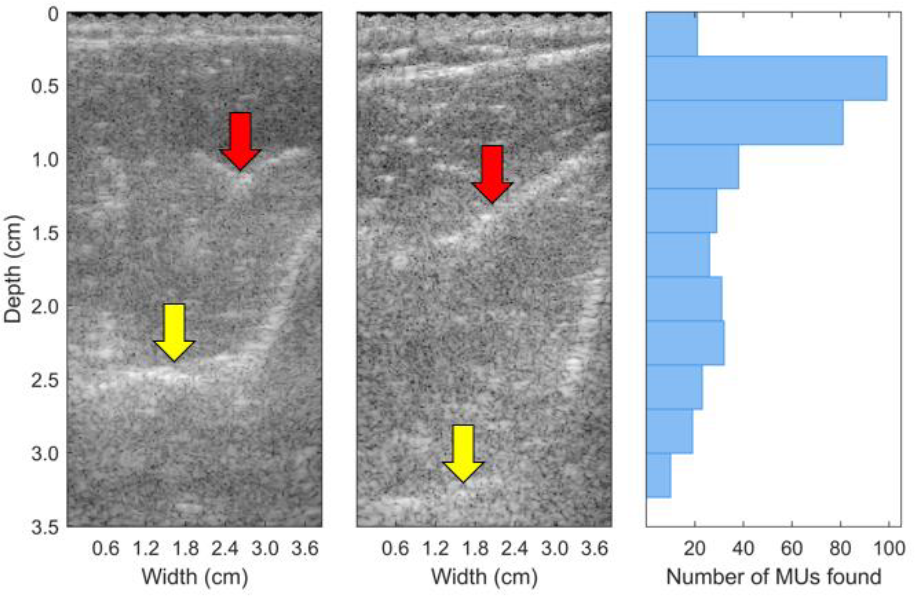
Left and middle: Example ultrasound (US) images from the tibialis anterior muscle of two participants, showing the end of the muscle (yellow arrow) and the location of the central aponeurosis (red allow). Right: The density of motor units (MUs) identified using US decomposition as a function of depth within the muscle. The highest density of MUs is identified in the 3-9 mm region, partially due to experimental bias (the use of online feedback based on stable recruitment of EMG detectable, and therefore superficial, units) and partly as all participants have muscle here. The dip at approximately 1 cm is due to the central aponeurosis, and the distribution tails off after 2.5 cm as some participants muscles end here (e.g., left US image).

In previous decompositions of US data, the RoA with respect to EMG was calculated using a window of 30 ms (± 15 ms around the EMG spike) [15], [21], which is extremely large. In contrast, we used a window of 0-5 ms following the EMG-detected spike after signal alignment. The time shift applied to the US signal represents the time between the EMG and US discharge times and likely reflects a combination of the excitation-contraction coupling and the identification of the ‘onset’ of the signal using this method. However, once this offset was removed, high RoAs are achieved in a small window, and variability was very low. Hence, although there was a small offset between the signals, the offset was relatively constant throughout the recording.

This work has shown for the first time that decomposition of US image series into individual MU discharge times, and therefore into the direct neural drive to the muscle, is possible across the whole muscle cross-section. While the results are already substantially superior to previous approaches, the method can be further refined. Firstly, whilst the validation provided in this paper used the same parameters, including window size, noise thresholding, and unit selection criteria for all participants, results could likely be improved with participant-wise adaptive selection of parameters. This would mirror the editing process routinely performed by experts on HDsEMG based decomposition [34]. Hence, work should be done into auto-selection of the best parameter space for a given participant. Furthermore, besides the inability to detect deep MUs, a key issue with EMG is its inability to decompose MU activity in dynamic contractions. Future work should test US decomposition on dynamic contractions alongside higher force-level contractions. Finally, as discussed above, recent work has shown some non-linearity in the summation of twitch velocities across MUs [39]. Although the method of linear convolutive decomposition presented here has shown a marked improvement in spike train estimation with respect to the linear instantaneous decomposition method, in the future, considering a non-linear approach, or introduction of nonlinear steps into this approach, may further improve the decomposition performance.

## V. CONCLUSION

In this work, we have presented the first methodology for full decomposition of the series of MU discharge times across an entire muscle cross-section, using ultrafast US. This has significance both for studying human neurophysiology and for non-invasive human-machine interfacing. It enables precise localization and further spatial information for MUs, as well as precise temporal information for times of APs and, therefore, spinal MN activity. We have provided an experimental validation for the proposed methodology. In contrast with commonly used sEMG, activity from deeper units can be decomposed, and further enriching information can be obtained about unit distribution, shape, and size. In contrast with other US decomposition techniques that cannot extract neural discharge times across the US image series, our method provides a reliable source estimation with accurate neural discharge times.

