## Supplementary Material for "Accurate Identification of Motoneuron Discharges from Ultrasound Images Across the Full Muscle Cross-Section"

### Supplementary Materials

(i) Non-MU US outputs

Fig. (1) shows an example of a non-MU output from the convolutive BSS of the US image series caused by the regular pulsations of a blood vessel in the imaging plane.

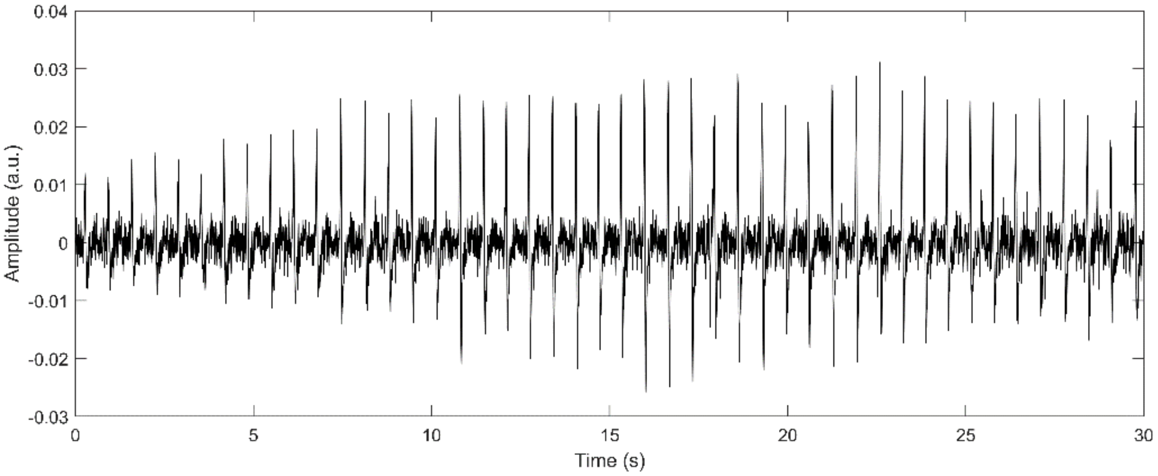


Fig. 1. Example non-MU output from the convolutive BSS attributed to the localised pulsations of a blood vessel. Peaks can be seen at an approximate frequency of 1.5 Hz, within the expected range for pulse rate.

(ii) Deep MUs detected by sEMG

In some cases, motor units MUs detected by both EMG and US are deeper than would be expected from the detection volume of sEMG (< 1 cm). One possible explanation of this is that the territory of the MU may be split, thus electrical activity may be distributed across the muscle cross section, with some electrical activity within the top 1 cm and thus detectable from the EMG.

To analyse this, we used the US detected discharge times to perform a spike-triggered average (STA) on the US derived velocity map series, using the methods described in [1]. Examples for two MUs which are detected by both EMG and US but are deeper than expected are shown in Fig. 3. In each case, regions which move synchronously with the MU are seen in the top 1 cm. It is therefore possible that these represent split territories and there is both motion and electrical activity in the top 1 cm which EMG has detected.

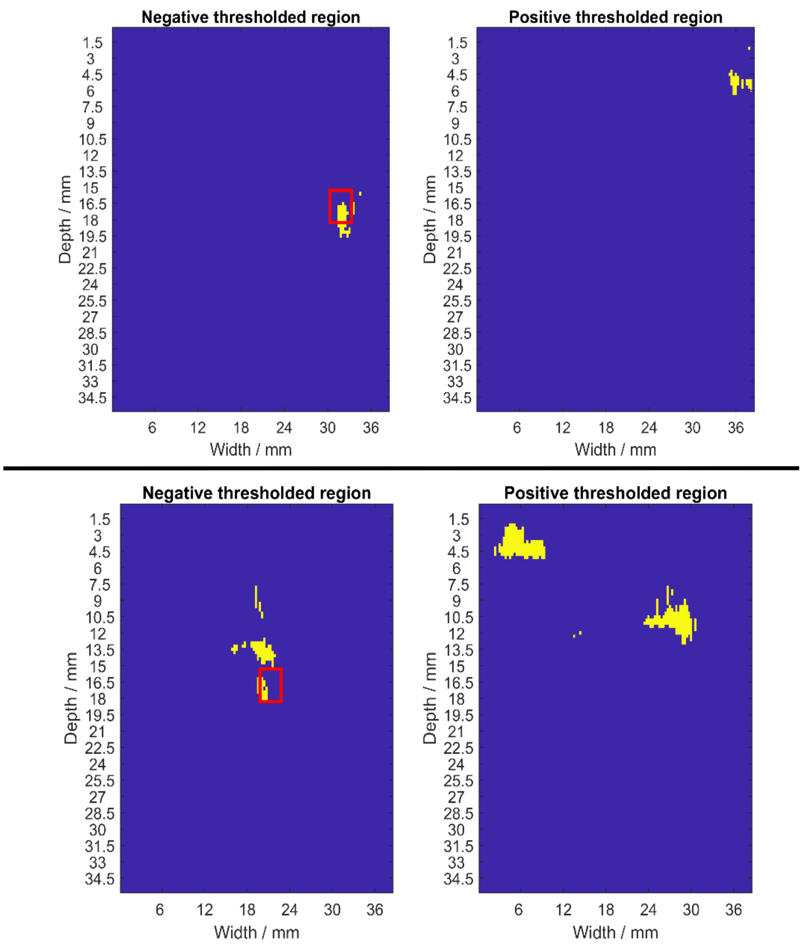


Fig. 3. Example spike-triggered average of velocity map series using ultrasound (US) detected discharge times. The US subgrid where the motor unit (MU) is detected is shown using a red box. Although in both cases the MU location is below the top cm of tissue, and therefore the unit should not be detectable using EMG, there is motion due to the MU in the top 1 cm of muscle suggesting that the territory is split.

[1] E. Lubel, B. Grandi Sgambato, D. Y. Barsakcioglu, J. Ibáñez, M. X. Tang, and D. Farina, "Kinematics of individual muscle units in natural contractions measured in vivo using ultrafast ultrasound," *J. Neural Eng.*, vol. 19, no. 5, 2022, doi: 10.1088/1741-2552/ac8c6c.
